# Insights into the microbial strain mediated impact on pest insect development

**DOI:** 10.1101/2022.05.03.490458

**Authors:** Kiran Gurung, Joana Falcão Salles, Bregje Wertheim

## Abstract

Molecular analyses of host-associated microorganisms have demonstrated the essential role that the microbiome plays in host development. Approaches targeting the sequencing of ribosomal genes have successfully identified key species of the host-associated microbiome. However, it remains unclear to what extent the strain-specific characteristics influence the outcome of the host-microbiome interactions. This is particularly important for insect pests, where microbial species might be used as targets for biocontrol purposes. Understanding strain-level variation represents thus a crucial step in determining the microbial impact on hosts. To investigate the microbial strain-level effects on an invasive insect pest, *Drosophila suzukii*, we compared the impact of monocultures and cocultures of different bacterial and yeast strains. We investigated whether different strains of *Gluconobacter* and *Pichia* differentially influenced the larval development of the pest. Fly trait measurements demonstrated beneficial, although variable, impact of these microbial strains on the fitness of *suzukii*. Using cocultures of microbial strains, we found that in some combinations, the beneficial effects were intermediate between those of the respective monocultures. In contrast, in other cases, strong inhibitory effects were observed. Hence, our study reports that strain-level effects within species are present in *D. suzukii*, reinforcing the importance of assessing the impact of associated microbiota on pest insects at the strain level.

**Highlights:**

- Microbial strains make up an essential part of the diversity of an insect host’s
- Characterizing and accounting for strain-specific impact on a pest’s life-history traits and different combinations of strains constitute an important step in our understanding of the pest management strategies.
- We investigated whether there was any strain-specific impact of bacteria and yeasts on the larval development of a frugivorous pest.
- We observed that strains varied in their impact, both as monocultures and cocultures, indicating their importance in modifying the host ecology.
- Our study adds to the growing literature on the importance of strains in pest insects.

## Introduction

Pest insects are generally associated with a myriad of metabolic and genetically diverse microbial species. However, studying the microbiota at species-level resolution is often insufficient as strains from the same species may differ largely in phenotypic traits, impacting their host associations (Barber et al., 2016; van Rossum et al., 2020). In addition, numerous factors determine and shape the strain diversity of their insect hosts, including genetic variations, metabolic capabilities, host-microbe coevolution and microbe-microbe interactions (Joop and Vilcinskas, 2006; Miraldo and Duplouy, 2019, Gasmi et al., 2021).

Diverse microbial strains reportedly structure the microbial communities and, eventually, their host’s ecology (Xie et al., 2019; Yan et al., 2020). Strains that belong to the same microbial species may differ substantially in the impact they have on their insect hosts, and they may affect multiple traits simultaneously. For example, numerous strains of *Pseudomonas syringae* had highly variable effects on the survival of the aphids and whiteflies following their infection (Smee et al., 2017). Strains of *Spiroplasma* differentially affected the fitness of aphid hosts in terms of offspring production as well as pathogenic resistance (Pang et al., 2018; McLean et al., 2020). McLean et al. (2020) demonstrated that a few *Spiroplasma* strains did not affect fungal sporulation, whereas others decreased sporulation. Another study involving five strains of *Pseudomonas chlororaphis* showed a differential effect on insect survival (Vesga et al., 2021). Furthermore, the microbial strains can also aid the host insect to cope or survive in challenging or new environments (Sugio et al., 2015; Pang et al., 2018), as witnessed in the case of the facultative endosymbiont *Arsenophonus* that influenced insecticide detoxification in a pest insect (Pang et al. 2018). These studies have highlighted the significance of understanding the microbial strain-level impact on insect pests.

In addition to the insect-microbe interactions, inter-strain microbial interactions (in cases of multiple co-infections) can potentially mediate the pest’s life cycle. Both *in vitro* and *in vivo* studies have demonstrated the importance of microbe-microbe interactions as major drivers in community dynamics and host life-history (Seth and Taga, 2014; Braga et al., 2016; Kroll et al., 2017; Venturelli et al., 2018). Specifically, multiple co-infecting microbial strains exhibit varied effects on the host’s fitness, depending on the types of co-infecting strains and their interactions (Narita et al., 2007; Kinnula et al., 2017; Mc Lean et al., 2018). When these strains compete for host resources, their interference may hamper host fitness, while facilitation among microbial strains could even enhance their beneficial impact on host fitness.

One member of the drosophilid family has evolved into a highly invasive pest, *Drosophila suzukii.* This insect has its origin in south and eastern Asia and has successfully invaded several regions of America and Europe (Walsh et al., 2011; CABI, 2021). More recently, *D. suzukii* has been reported from the berry growing region of sub-Saharan Africa (Kwadha et al., 2021). Cherry, strawberry, raspberry, blackberry, blueberry, and grapes are among some of the preferred fruits, regularly oviposited by these flies (Hausier et al., 2011). Additionally, other fruit crops and potentially ornamental plants are reported to be susceptible to the attack of *D. suzukii* (Walsh et al., 2011; Kenis et al., 2016). Females lay eggs into the ripening fruits by piercing the fruit skins using their serrated ovipositor. This infestation may lead to the rotting of the fruits, making this insect solely responsible for large commercial loss in the infested regions (Walsh et al., 2011).

The microbial communities of this pest fly are highly variable in composition and diversity, both for flies and the infested fruits, with key microbial species belonging to the bacterial families Acetobacteriaceae and Enterobacteriaceae and the fungal phylum Ascomycetales, to name a few (Bing et al., 2018; Fountain et al., 2018; Solomon et al., 2019; Gurung et al., 2022). Furthermore, a handful of studies have verified the importance of the microbiome during the development of *D. suzukii*. For instance, the incorporation of live and killed microbes into the diet of the flies has shown that several bacterial and yeast taxa are important already at the juvenile stage (Bing et al., 2018; Hamby et al., 2012). Moreover, bacterial and yeast strains impact the fly’s life-history traits, such as larval survival, adult longevity, adult body size and oviposition (Mazzetto et al., 2016; Lewis and Hamby, 2019; Solomon et al., 2019; Spitaler et al., 2020). Therefore, understanding the interactions and functional roles that microbes might have in the life history of invasive pests such as *D. suzukii* is an important aspect of pest management. Overall, these studies demonstrate that bacteria and yeast contribute in shaping the life-history traits and also draws further focus on their roles at the strain level. Integrating strain-level studies might also explain the variations observed when studying the pest’s microbial ecology.

In this study, we asked whether strain-specific impact could be observed in the juvenile development of the *D. suzukii*. We were also interested in understanding whether and how microbe-microbe interaction would potentially affect this development. Therefore, we investigated whether the impacts of these strains on host development vary, both when they are assessed individually and when assessed in different combinations. We addressed these questions using two bacterial strains of *Gluconobacter cerinus* and three yeast strains, two of which belonged to *Pichia kluyveri* and one strain belonging to *Pichia terricola*. All microbial strains were isolated from wild-caught *D. suzukii* in the Netherlands. We first profiled these microbial strains to assess their substrate utilization patterns and to verify the existence of phenotypic differences among these strains. Next, by adding monocultures and cocultures of the bacteria and yeasts to the juvenile diet, we monitored their impact on the overall development of the larvae by measuring fitness traits such as pupal and adult emergence and developmental time.

## Results

### Characterization of individual bacterial and yeast strains

#### Bacteria

We profiled the metabolic capability of the two bacterial strains of *Gluconobacter cerinus*, GC1 and GC2, using Biolog plates-GENIII and assessed their use of functional groups in terms of average well colour development (AWCD). The Biolog-GENIII provides insight into the metabolic capability of aerobic bacteria and contains multiple wells with embedded substrates belonging to different functional groups, namely sugar, sugar alcohol, amino acids, hexose phosphates, carboxylic acids, hexose acids and inhibitory salts. Strain GC2 showed a more diverse profile than GC1 (figure 1a). We observed a significant difference in the utilization pattern of the different functional groups according to the average well color development (Kruskal Wallis test, *X^2^*=55.36, p-value<0.001). Sugar alcohols were among the highest metabolized functional groups by GC1, whereas GC2 variably used all functional groups and developed in the presence of salts. Almost all functional groups contributed to the differences in the utilization patterns of the two strains (Dunn test, p-value <0.05).

**Fig. 1.**
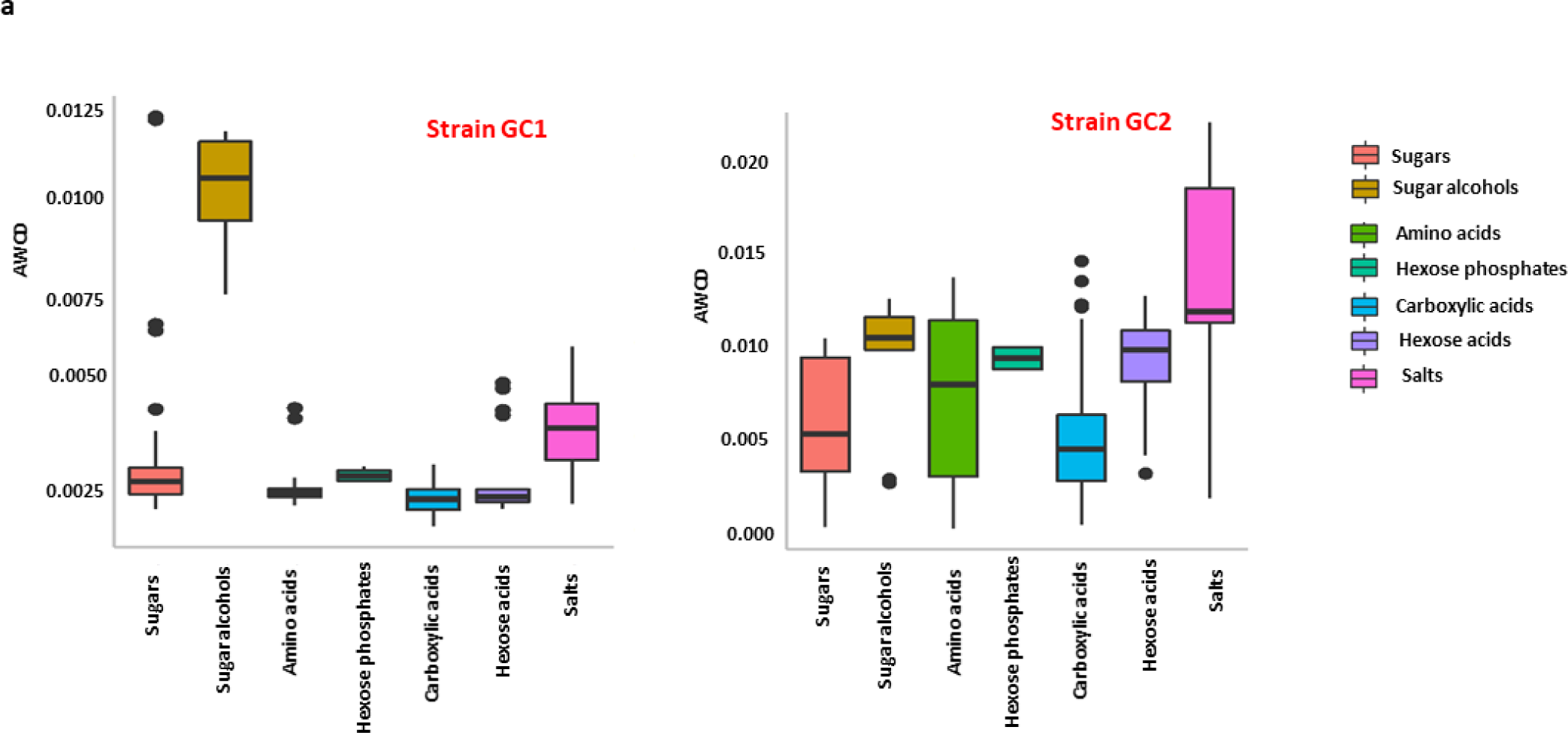
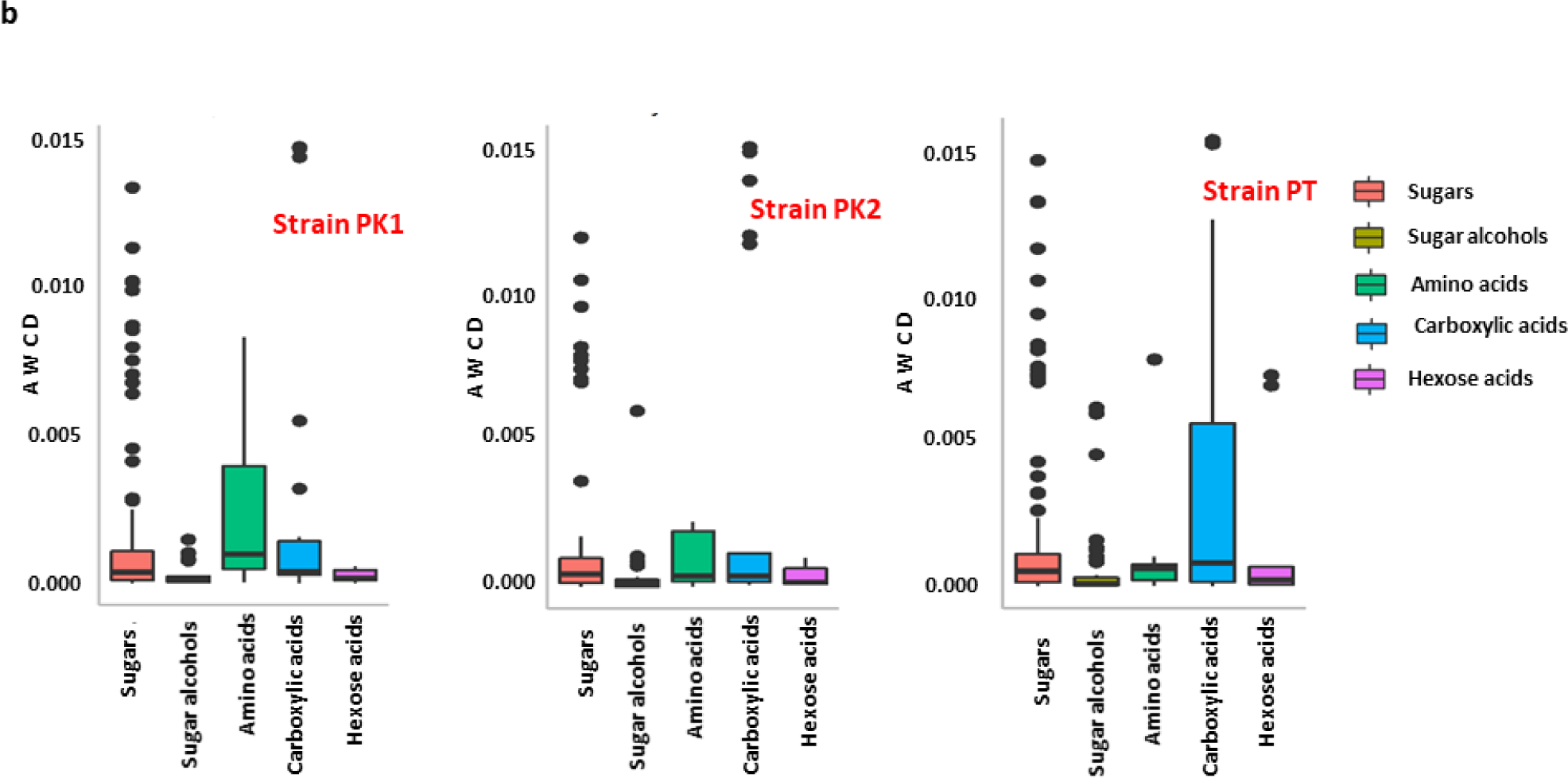
Substrate utilization profile of bacteria and yeast strains over major functional carbon groups, calculated as average well colour development (AWCD) 70h after inoculation. **(a)** AWCD of bacterial strains of *Gluconobacter cerinus* GC1 and GC2 was determined using BIOLOG plates GENIII and varied across the functional groups (Kruskal-Wallis test *X^2^*=55.36, p-value < 0.001). **(b)** AWCD of three yeast strains, two of *Pichia kluyveri* (PK1, PK2) and one of *P. terricola* (PT), was determined using Biolog plates YT plates for yeast strains and differed across functional (Kruskal-Wallis, test *X^2^*=50.84, p-value < 0.001).

### Yeasts

We profiled the metabolic capability of three yeast strains, two of *Pichia kluyveri* and one of *P. terricola*, PK1, PK2 and PT, using Biolog plates-YT. Biolog-YT plates contain substrates belonging to functional groups sugar, sugar alcohols, amino acids, carboxylic acids, and hexose acids and provide the metabolic potential of (aerobic) yeasts. Functional group utilization across the yeast strains was variable. And we observed significant differences in the utilization of the functional groups (fig. 1c, Kruskal-Wallis test, *X^2^*=50.84, p-value < 0.001), with carboxylic acids, hexose acids and amino acids contributing to the difference between groups (Dunn test, p-value <0.05). While strain PT’s metabolic capacity for the carboxylic acids was the highest, the metabolic capacity for the amino acid groups was the highest for strain PK1.

## SURVIVAL

### Bacteria

The strains showed varied effect on the survival of eggs to the pupal stage (Fig. 2a & 2b; GLMM, binomial: p-value < 0.001). Survival up to the pupal stage for juveniles developing in the presence of a monoculture of GC1 was higher than the ones in the monoculture of GC2 (Tukey test, p-value <0.001) and also the coculture (Tukey test, p-value<0.001). Survival was not significantly different between GC2 and coculture (Tukey test, p-value = 0.458). Pupal survival in the bacterial treatments was still lower than when compared to the HKY food (Tukey test, p-value <0.001), while for the NYC plates, i.e. the control comprising minimal food without any microbial additions, survival was significantly lower than in the all the other treatments (Tukey test, p-value < 0.001).

**Fig. 2.**
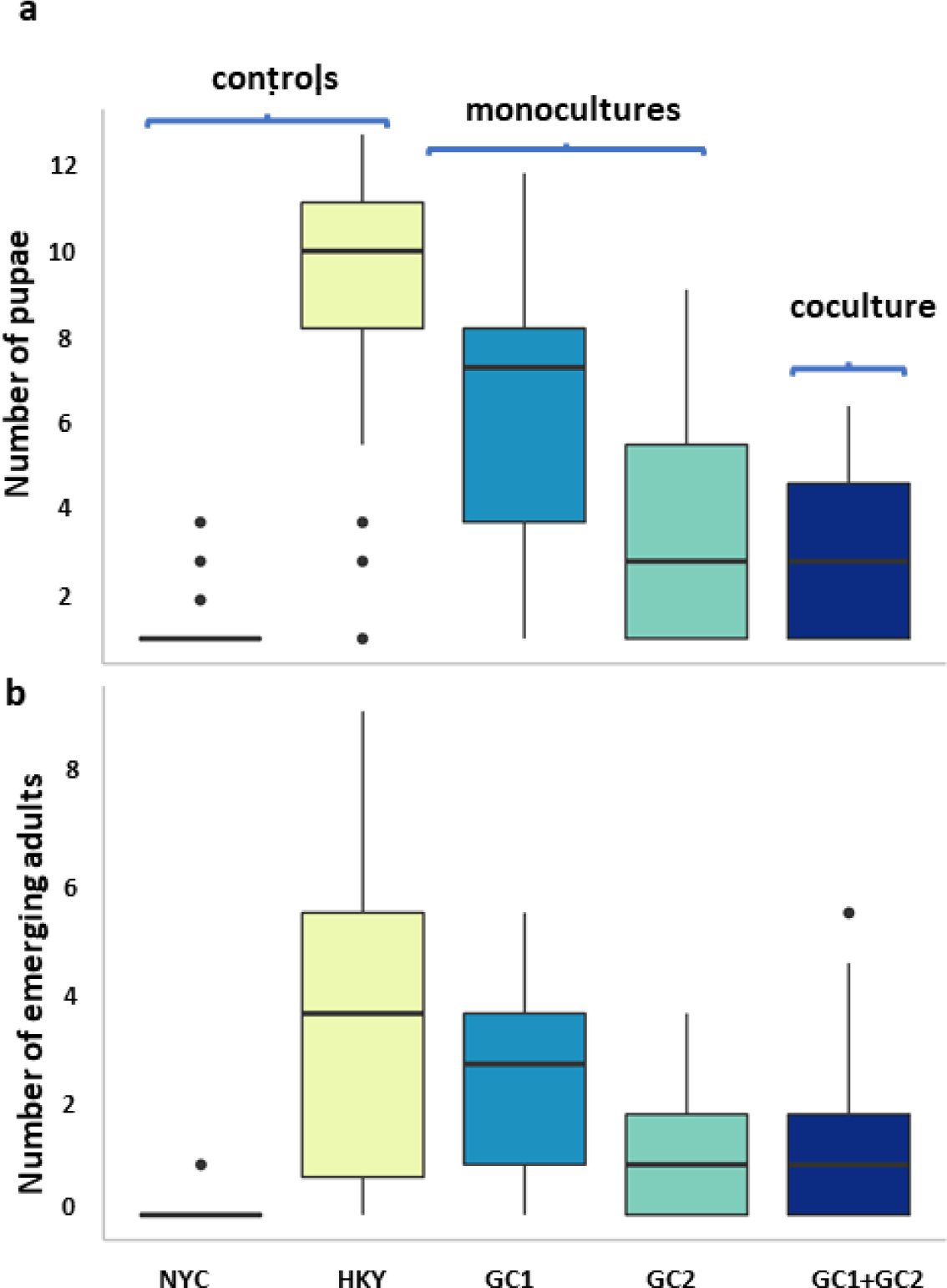
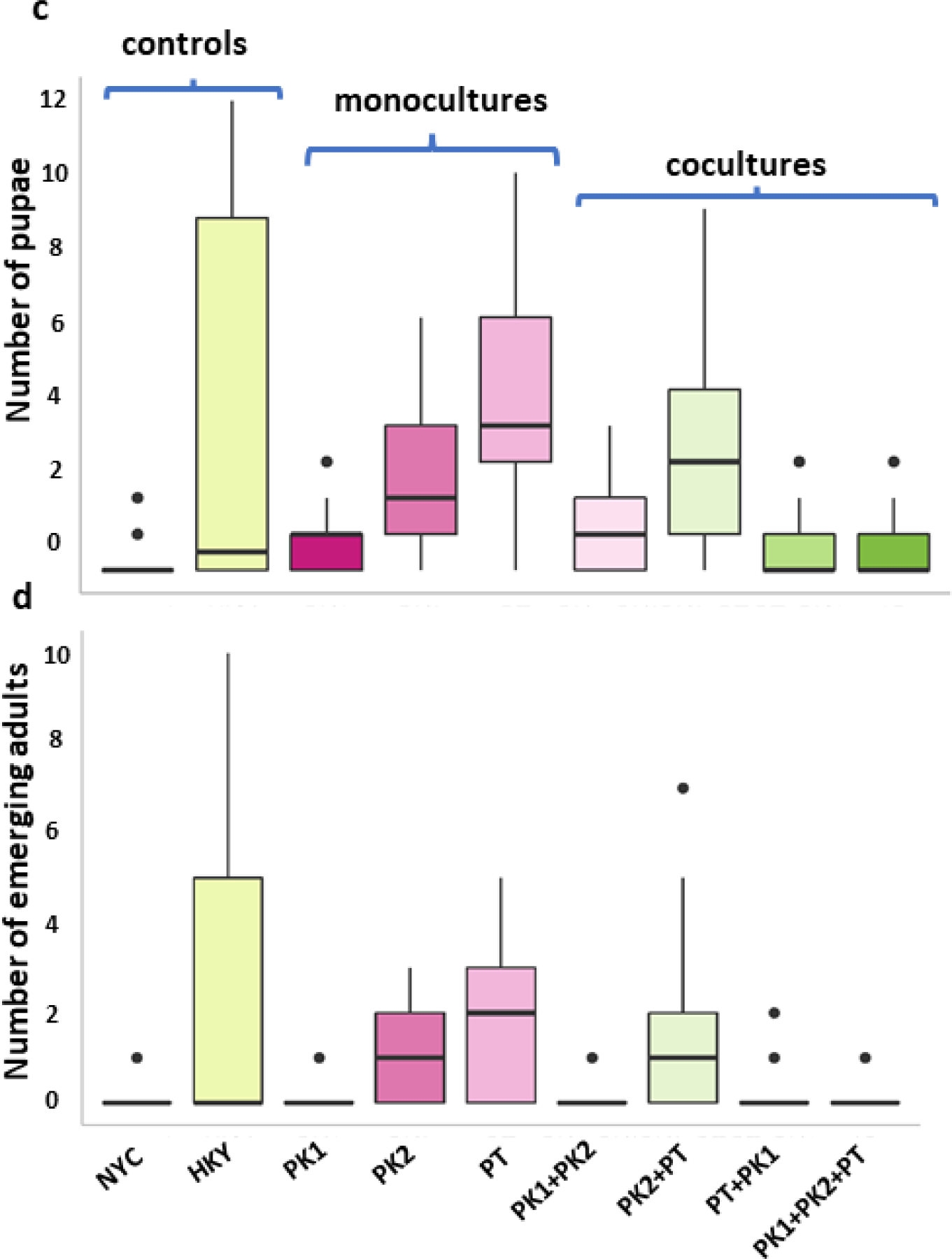
Egg-to-pupal and egg-to-adult survival of *D. suzukii* in the presence of different microbial strains. The number of surviving a) pupae and b) adults (out of 13 eggs that were added to each plate) differed significantly across monocultures and cocultures of two bacterial strains of *G. cerinus* (GC1 and GC2) (GLMM: p-value < 0.001). Significant differences were also noted for c) pupal and d) adult survival with monocultures and cocultures of three yeast strains, two of *P. kluyveri* (PK1, PK2) and one of *P. terricola* (PT) (GLMM: p-value < 0.001). [NYC: No yeast control, HKY: Heat killed yeast control]

Also, the survival to adult stage differed significantly across monocultures and cocultures (GLMM, binomial: p-value < 0.001), with significant differences between GC1 and GC2 (Tukey test, p-value < 0.001), but no difference between each of the monoculture GC2 and the coculture (Tukey test, p-value = 0.450). Survival in the adult stage in the HKY diet was significantly higher than in all other diets (Tukey test, p-value < 0.001).

#### Yeasts

There was a significant difference in the survival up to the pupal stage for the different yeast strains for both monocultures and cocultures (fig 2c & 2d, GLMM, binomial: p-value < 0.001). Survival of the juveniles up to the pupal stage was the highest in the diet supplemented with a monoculture of the PT strain, compared to the two other monocultures. Among cocultures, we also observed significant differences (Tukey test, p-value < 0.001), with a rather striking pattern: any coculture containing PK1 would result in equally low survival, while the coculture of PK2+PT gave a survival that was intermediate to the two monocultures PK2 and PT. Survival for the diet with the coculture of all three strains (PK1+PK2+PT) showed a significant difference with other treatments (Tukey test, p-value < 0.001), except for PK1, PK1+PK2, and PT+PK1, respectively (Tukey test, p-value = 0.906, p-value = 0.859, p-value = 0.96). Survival in the HKY diet was also higher when compared to other yeast treatments but did not differ significantly from the PT and PK2+PT (Tukey test, p-value = 0.320, p-value = 0.878).

Up to the adult stage, survival was highest in monocultures PK2, PT and coculture PK2+PT, while we observed stark reduction in fly survival until adulthood for monoculture PK1 or cocultures with PK1. Overall, we observed a significant difference in the adult emergence across all the treatments (GLMM, binomial: p-value < 0.001). For survival up to adult emergence, among the monocultures, pairwise contrasts showed significant differences across all the strains, with the highest survival on the PT strain (Tukey test, p-value < 0.001). All three monocultures showed significant differences by pairwise comparison (Tukey test, p-value = 0.005, p-value < 0.0001). For cocultures, no significant difference was observed between PK1+PK2 and PT+PK1 (Tukey test, p-value = 0.655). Among those treatments that showed inhibitory effect on the adult emergence, PK1+PK2, PT+PK1, and PK1 showed no significant difference compared to the coculture PK1+PK2+PT (Tukey test, p-value = 0.655, p-value = 1, p-value = 0.537). The adult emergence on HKY food was significantly higher than all other treatments, except when compared to the monoculture PT (Tukey test, p-value = 0.991). Adult emergence in the NYC minimal diet was low and comparable to those on PK1, PK1+PK2, PT+PK1(Tukey test, p-value = 0.444, p-value = 0.543, p-value = 1). We also observed that adult emergence in the HKY diet was more variable in the yeast experimental set-up when compared to the set-up comprising bacterial strains.

### Developmental time

#### Bacteria

Developmental time to the pupal stage and adult emergence were the shortest in the control diet with heat-killed yeasts (HKY). Adult emergence time in the case of the minimal diet without microbial strains (NYC) was longer. Strain GC2 exhibited a longer duration for the adults to emerge than GC1. The mean developmental time (measured in the number of days) between the juvenile stage and pupation ranged from 12-18 days for monocultures and cocultures and 17 days for the NYC minimal medium, while the egg-to-adult developmental time spanned a total duration of 21-26 days (fig. 3a & 3b). The coculture showed a shorter developmental period when compared to the other two monocultures. The rich medium HKY showed a shorter mean developmental time of 9.3 and 15.14 days. This developmental time differed significantly across the treatments (GLMM, Poisson: p-value < 0.001). The coculture and the rich food HKY contributed to this difference (Tukey test, p-value < 0.001).

**Fig. 3.**
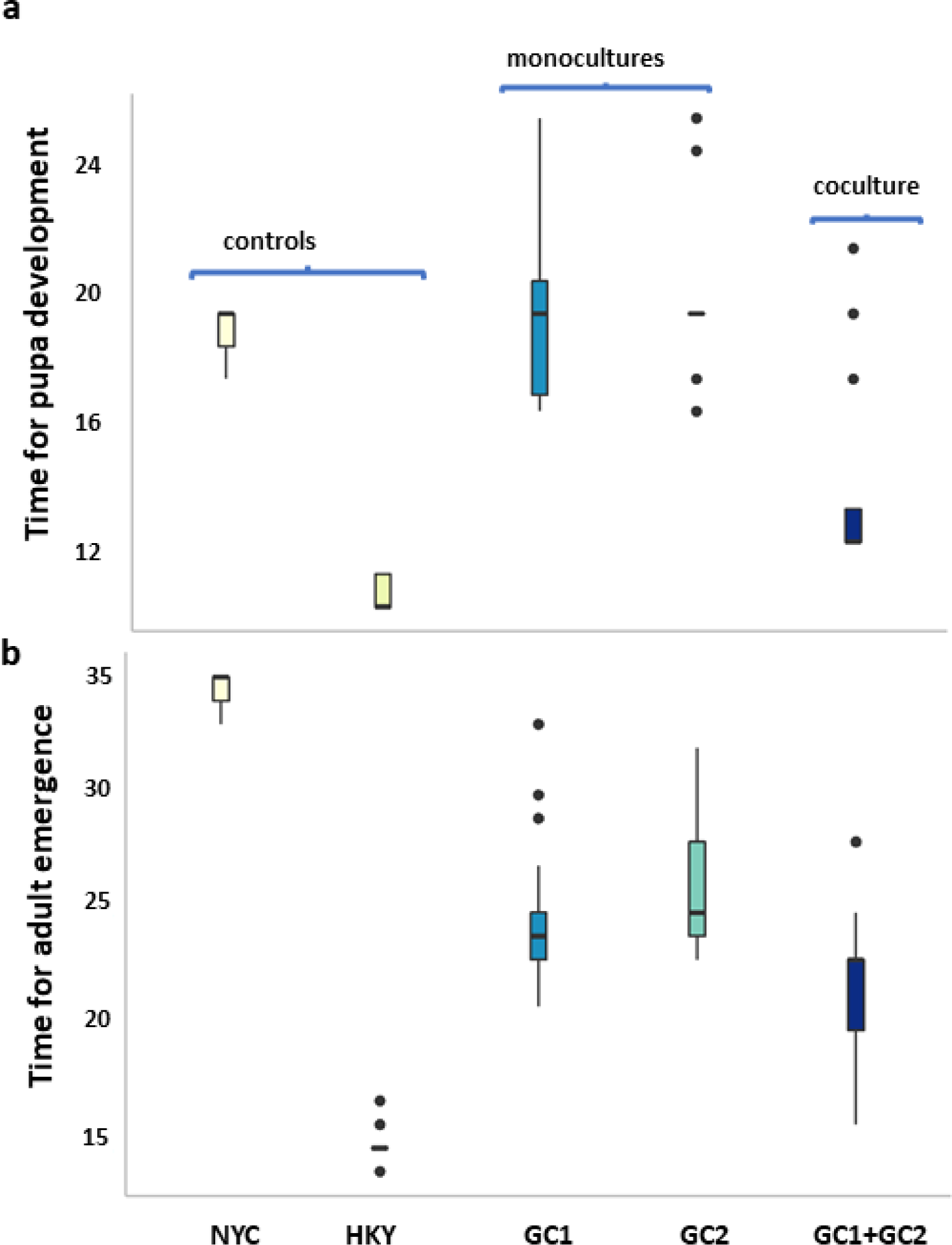
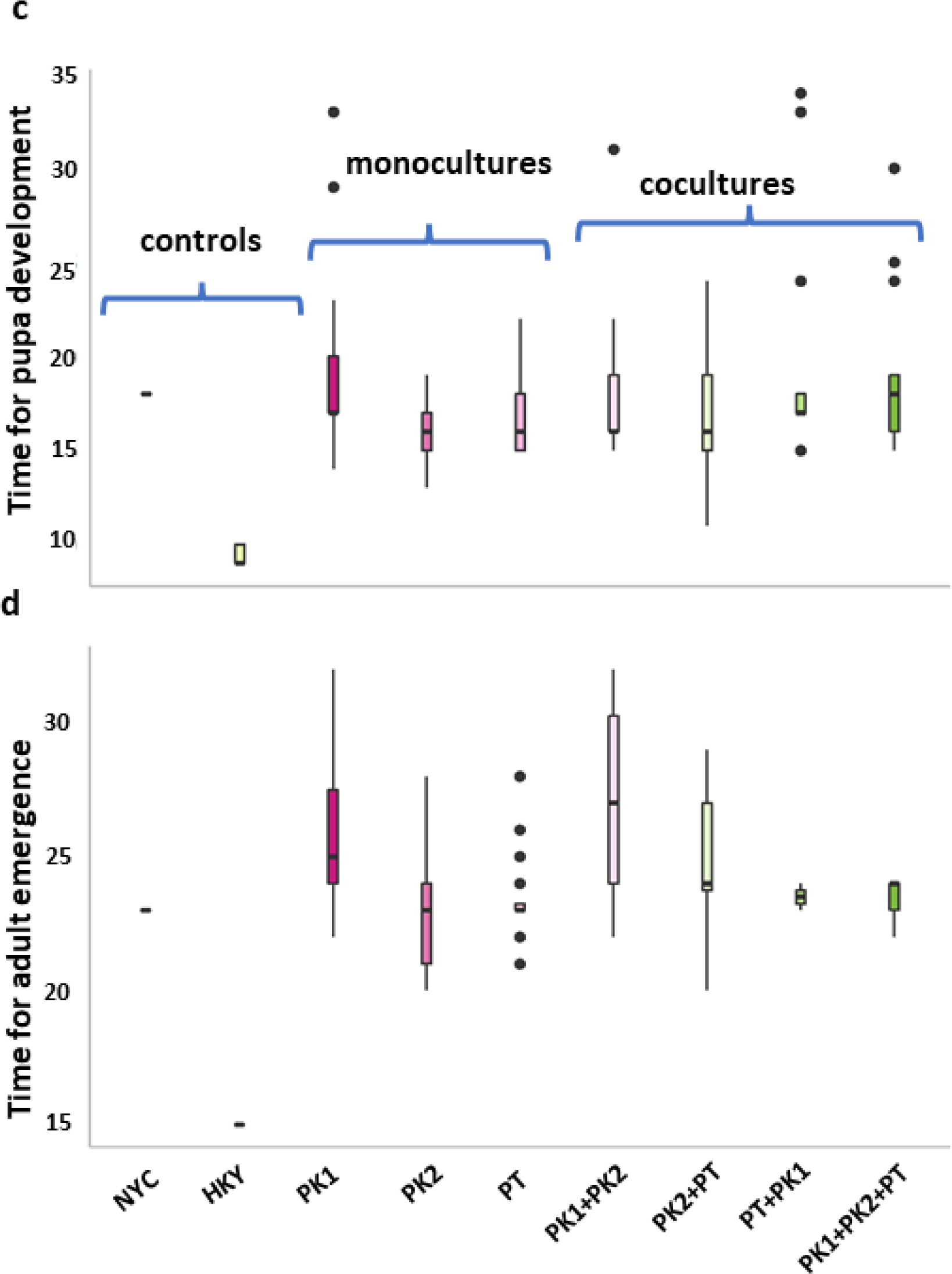
Developmental time (in days) for the *D. suzukii* eggs to reach pupal and eventually adult stage in the presence of different microbial strains. The data presented here and the analysis conducted in the dataset with non-zero values. For bacteria, significant differences existed across the developmental time to reach a) pupal stage and b) adult stage (GLMM: Poisson p-value < 0.001). For yeasts, we observed a significant difference in reaching c) pupal and d) adult stage (GLMM: Poisson p-value < 0.001). [NYC: No yeast control, HKY: Heat killed yeast control]

#### Yeasts

In most cases, pupation required 16-19 days and adult emergence required about 23-27 days. There was a significant difference in the developmental time of egg-to-pupa and egg-to-adults across monocultures and cocultures of yeast strains (fig. 3c & 3d, GLMM, Poisson: p-value < 0.001). The pairwise comparisons indicate that this reflects the differences between the HKY and the NYC diet with each of the other treatments (fig. 3c & 3d). We observed no significant difference in pairwise comparisons among the other yeast monocultures and cocultures. For the food with HKY, the mean developmental time for egg-to-pupal and egg-to-adult stages was 9.2 days and 15 days, respectively. This was significantly shorter than in any of the other treatments.

### Body size

#### Bacteria

The tibia length (used as a proxy for body size) differed significantly across all the treatments (measured for a subset of 29 flies/treatment), with 39.4% of variance explained by different treatments and 11.1% explained by the sex of the flies (fig. 4a, mixed-model ANOVA, p-value < 0.001). According to the post-hoc test, the rich HKY diet resulted in larger tibia length for both males and females than the monocultures or coculture (Tukey test, p-value < 0.0001).

**Fig. 4.**
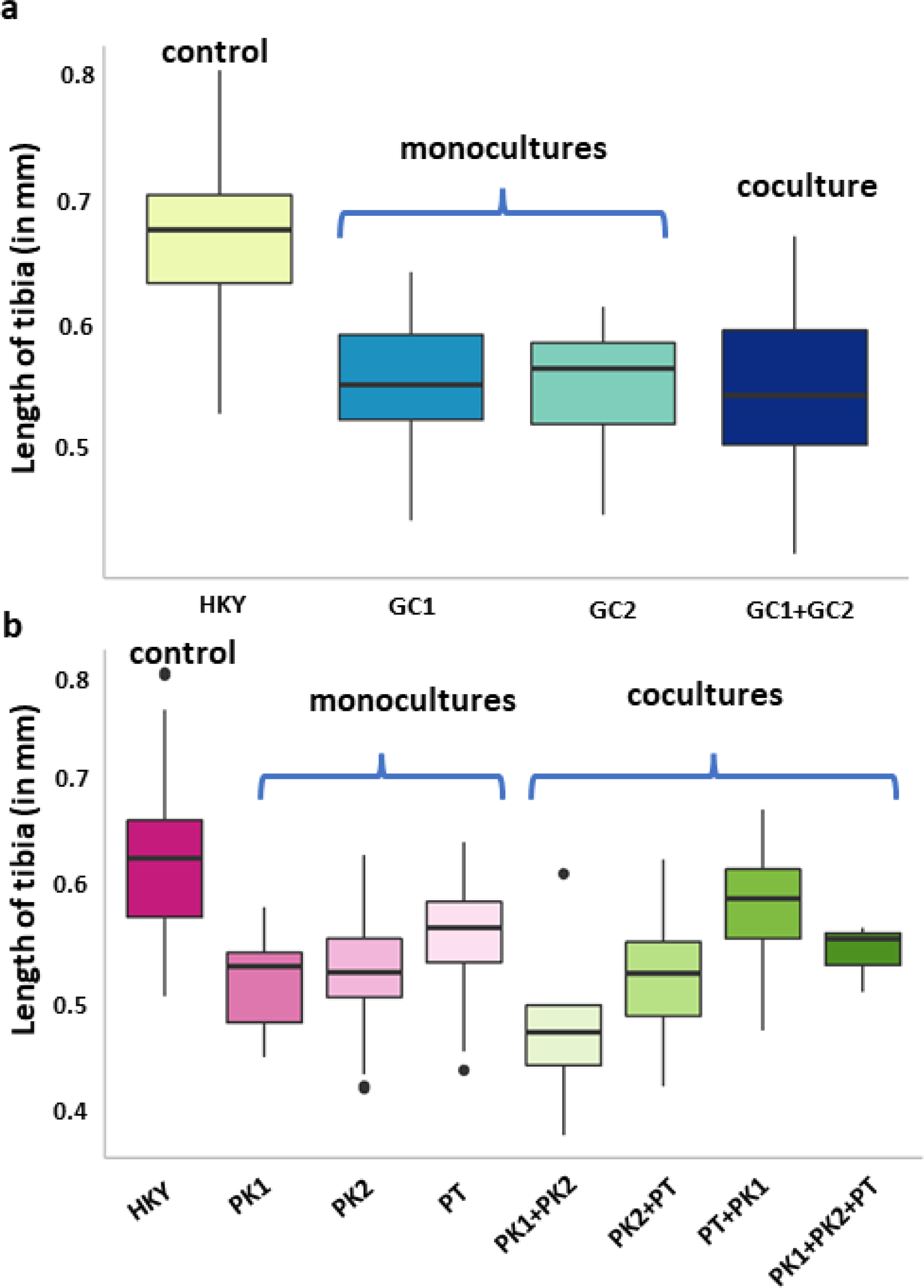
Tibia length as a proxy for *D. suzukii* adult body size for juveniles developed on different microbial strains. **(a)** For bacterial strains, tibia length differed significantly (ANOVA, p-value < 0.001), with adults emerging from the rich diet HKY being larger than those developing on either of the two *G. cerinus* strains (GC1/GC2) **(b)** For yeast strains, a significant difference was noted (p-value < 0.001) for sample sizes ranging from 5 flies to 30 flies. However, analysis of a subset of 9-11 flies showed a significant difference in tibia length, which was solely contributed by adults from rich diet HKY (p-value < 0.001)

#### Yeasts

Tibia length differed significantly across different treatments (for a subset of 9-11 flies/treatment) (fig. 4b, mixed-model ANOVA, p-value <0.001), with 22% of the variance explained by the treatment and 4% by the sex. The significant differences were in the tibia length of the flies for both sexes from the rich medium HKY compared to the monocultures and cocultures (Tukey test, p-value < 0.0001).

## Discussion

In our study, we found empirical evidence for the importance of microbial strain-level variation in the life history of a pest insect. The supplementation of minimal rearing substrates with different bacterial and yeast strains had variable fitness consequences for *D. suzukii* juveniles in terms of larval and pupal survival, developmental rate and growth. We tested microbial strains belonging to *Gluconobacter* and *Pichia*, which are commonly associated with the niche of *D. suzukii* and even other frugivorous insects (Hamby et al., 2012; He et al., 2017). These microbial strains used in our study differed phenotypically, as evidenced by differential substrate utilization patterns (according to their average well color development). The survival of juvenile *D. suzukii* from eggs to the pupal and adult stage differed significantly when reared on monocultures and cocultures of two bacterial strains of *G. cerinus*. Also, across different yeast monocultures and cocultures, we observed significant differences in the surviving number of pupae and adults. Here, one of the monocultures resulted in high survival rates, whereas another strain nearly always precluded any survival of the juveniles, both when in monoculture and cocultures. For the developmental time, a similar pattern of significant differences existed across our treatments of both bacteria and yeasts. The body size of the emerging adults was not significantly different across the microbial strains but was smaller compared to juveniles reared on rich food containing heat-killed yeast. However, it should be noted that only a small number of eggs developed to the pupal or adult stage in some treatments, which could have contributed to the non-significance of body size. All the fly assays aligned largely: microbial strains with lower survival also yielded slower development. Furthermore, the inhibitory effect posed by one of the yeast monocultures on the juvenile’s survival, developmental rate and growth were also dominant when mixed in cocultures. Our findings thus highlight that these microbial strains vary in how they influence the pest’s development in terms of their survival, both in monocultures and in cocultures, where microbe-microbe interactions may also play a role. We still have limited knowledge of mechanisms that tie the idea of microbe-microbe interactions to pest fitness. However, in *D. melanogaster*, specific characteristics of individual taxa shaped the fly fitness, fecundity and developmental time, with both positive and negative potential impacts on the host (Gould et al., 2018).

Both for the bacterial and the yeast strains, the effects of cocultures on fly survival, developmental rate and growth were not simply additive or intermediate between the impact of monocultures. For example, PT could not overcome the strong negative effect caused by one of the other two strains in cocultures, PK1. Although, as a monoculture, PT indeed showed high survival for the juveniles, it failed to enhance juvenile survival when it was combined with PK1 and when PK2 was added. To a lesser extent, this was also the case for the monocultures and cocultures of the bacterial strains, where the coculture resulted in a lower juvenile survival than one of the two monocultures. In another study, at an inter-kingdom level, Bing et al. (2018) showed that cocultures of yeast and bacteria influence the developmental time in *D. suzukii.* This emphasizes that we need to acknowledge the enormous diversity among different microbial strains the hosts are associated with and carefully address the potential interactions among these microbial strains that can result in unpredictable combinatorial effects.

Bacteria *Gluconobacter* belongs to the family of Acetobacteriaceae, which is highly prevalent in drosophilids (Crotti et al., 2010; Vacchini et al., 2017; Gurung et al., 2022). Members of the acetic acid bacteria (AAB) are the primary component of microbial communities in unrotten and rotten fruits (van Keer et al., 1981) and other environments rich in sugar, alcohol and acids. The AAB metabolize compounds belonging to the functional classes of alcohols and sugars (Raspor and Goranovič, 2008; Deppenmeier and Ehrenreich, 2009; Lynch et al., 2019). The genus *Gluconobacter* specifically utilizes sugars such as glucose to convert them into gluconic acid (Dwivedi, 2020). In a sucrose rich environment, this particular genus synthesizes levans from sucrose using the enzyme levansucrase (Jakob et al., 2019), enabling them to establish their niche in sugar-rich environments. In non-*Gluconobacter* spp., other variants of these enzymes are said to act as virulent factors against plants (Katzen et al., 1998). In our results, we observed that both bacterial strains use sucrose. Besides, although the substrate utilization pattern varied between the two bacterial strains, the sugar alcohols were consistently being used by these two strains (Deppenmeier & Ehrenreich, 2009; Matsushita et al., 2003). These bacterial genera tend to necrotize and rot fruits, a prominent example being pink rot disease in pineapples (Van Keer et al., 1981; Dwivedi, 2020). They usually grow at an optimal pH of 5-6, with few strains growing at pH as low as 3. *Gluconobacter cerinus* differs from most other bacterial species and even some AAB due to specific lipid composition and the presence of dehydrogenases engaging in carbohydrate metabolism (Tahara et al., 1976; De Muynck et al., 2007; Sainz et al., 2016). These strains have been routinely isolated from rotting samples of berries or pears and, due to their metabolic potential, have been assigned the primary mediators in fruit rots (Wieme et al., 2014; Sombolestani et al., 2021). They are also found in other frugivorous invasive insect pests and appear to induce rotting like symptoms in fruits (He et al., 2017). The study by He et al. (2017) demonstrates the pathogenic potential of *G. cerinus* CDF1, a strain intimately associated with a frugivorous invasive pest, *Bactrocera dorsalis*. However, the exact causal relationships between fruit rots and these bacteria remain to be confirmed.

The two *Gluconobacter* strains that we isolated from wild *D. suzukii* reared from infested cherries promoted larval survival as monocultures, compared to the microbe-free food source, although to a different extent for the two strains. This suggests that the microbial strains had a positive effect on the nutritional value of the food source. In addition, when they were combined in a coculture, the effect on the developmental time was less than intermediate for the two monocultures. Further, survival in presence of cocultures did not differ much from one of the monocultures. We do not have any clear explanation for these effects posed by the cocultures. Perhaps this might reflect the strain mediated inhibition that comes into force in coculture conditions, which primarily reflects the activity of individual strains. Alternatively, this effect could have been due to the lower initial density of one of the strains in co-cultures, which may be below the minimum threshold for microbial growth or beneficial effects for larval nutrition. Still, it seems plausible that the interactions between the microbial strains negatively affected their nutritional value to the larvae.

*Pichia* spp. are among the Ascomycetales involved in alcohol production. These yeast species assimilate glucose and produce a variety of aromatic compounds from ethanol and esters *in vitro* (Vicente et al., 2021). The utilization of carboxylic acids and amino acids by *P. kluyveri* strains was higher than the utilization of the other functional groups. In *P. terricola*, utilization of carboxylic acid was also the highest among the functional groups and even higher than in the other two *Pichia* strains. *Pichia* is regularly found in uninfested and infested fruits and insect guts and can influence the insect’s life history in several ways (Stefanini, 2018). This genus is also associated with *D. suzukii* (Hamby et al., 2012; Fountain et al., 2018). Other than their nutritional roles, these yeasts are also studied for their insect-luring capability (Lewis and Hamby, 2019; Lasa et al., 2019). While they support juvenile development, their role as luring agents is less pronounced than other yeast species (Lewis and Hamby, 2019; Lasa et al., 2019). Specifically, the species *P. kluyveri* has a variable effect on the survival of the *D. suzukii* flies (our work and Lewis and Hamby, 2019). Hence, we speculate that the variation of the two *P. kluyveri* strains in our study indicates the strain-specific impact on the insect life history of *D. suzukii.* Less is known about the species *P. terricola* in relation to insect microbiome studies. However, this species tends to boost alcohol and aroma production with its associated fruits (Liu et al., 2015; Vincente et al., 2021). In our experiment, the *P. terricola* strain had the most beneficial effect of the three yeast strains tested, both in survival, developmental time and body size of emerging adults. It should be noted that *Issatchenkia terricola* is thought to be synonymous with *P. terricola* (Vincente et al., 2021). Previously, *I. terricola* was reported to positively impact *D. suzukii* fitness (Lewis and Hamby, 2019). Nevertheless, we are unsure whether the two species are indeed synonymous. We found strain-specific differences for the other two yeast strains of *P. kluyveri* used in our study.

In addition to the surviving numbers of juvenile *D. suzukii* other fitness traits such as developmental time and body, size determines the success rate of pests (Hu et al., 2008; Shelly, 2018; Tran et al., 2018). A slow development rate or prolonged development time may place the larvae at a competitive disadvantage when the food source is ephemeral and may run out before all individuals complete development. Plus, shifts in developmental time might also affect the risk of predation or parasitization (Chown and Terblanche, 2006). In our study, we observed that, when survival was low, the few individuals who managed to reach the pupal or adult stage took substantially longer time, which was comparatively more prolonged than those emerging on the rich diet. Another important trait for fitness is body size, as it is positively correlated with fecundity in females and with mating success in males (Shelly, 2018; Tran et al., 2018). In our study, adults emerging from the rich diet containing the heat-killed yeasts were the biggest of all compared to the other treatments with live bacterial and yeast strains. We did not observe a strain-specific impact on the fly’s body size but were limited in sample size for some treatments due to low survival. While microbes and microbe-microbe interactions can influence a broad range of fitness traits in various other insects (Douglas, 2018; Gould et al., 2018; Lewis and Hamby, 2019), in our experimental set-up, the trait that was most prominently affected by the microbial strains was the survival of both pupae and adults.

Based on our studies, we cannot infer whether the differential impact of microbial strains was driven by direct or indirect interactions among the microbial strains, which still needs further testing. For example, because of different metabolic capabilities, two co-infecting strains might either construct their micro-niches or crossfeed through reciprocal exchange of metabolites and potentially demonstrate additive or synergistic effects on the infecting hosts (Hassani et al., 2018; Niu et al., 2020). In addition, inter-strain competition also results in an antagonistic effect where the antagonizing strain might play a crucial role in controlling the host (Russel et al., 2017; Hassani et al., 2018). Such interactions leverage several mechanisms, from producing signals engaged in quorum sensing to secondary metabolites (Braga et al., 2016).

Our study was performed in lab-based conditions, and we tried to control for confounding factors as much as possible. However, in the wild, many factors such as the interaction with other plants, insects and the host’s microbial community members could further influence the strain-specific impacts of microbes on insect fitness. Besides, the genotypic difference in flies also determines their success, and they can reportedly regulate the microbial community composition (Early et al., 2017; Lee et al., 2019). Hence, it should be noted that fly strains may also differ in how they respond to microbial supplementation in general and specific microbial strains. Nevertheless, our study highlights the significance of bacteria and yeasts at a strain level in determining the pests’ fitness.

## Significance

In our study, using culturable microbes, we found that different strains of bacteria and yeasts showed variable effects on the fitness of an insect host. Most strains improved the nutritional value of the larval feeding substrates, yet in some combinations, strong inhibitory effects seemed to surface in cocultures. Hence, our results confirmed a dependency of *D. suzukii* on microbes for larval development, while it also disclosed vulnerabilities. Furthermore, it emphasized the immense diversity among microbial strains in their effects on host insects and the complexity of strain-strain interactions. Understanding the mechanisms driving these microbial strain-level effects on insects could have implications for pest management.

## Limitations

The two main limitations of this study are as follows:

1 – In addition to larval development, oviposition as well as defense mechanisms against natural enemies are among some of the important fitness traits for a pest. Therefore, future work should also focus in scoring these traits. This would then broaden our understanding on whether (and to what extent) strain specific interactions shape the fitness traits altogether.

2 – Future studies based on *D. suzukii* microbiomes, could also focus on understanding similar interactions in wild conditions. Currently, we do not know to what extent strain specific interactions are in effect under semi-field and open field conditions. Hence, testing them in the wild, to some extent, will guide our evaluation of the differences between the lab-reared and wild flies.

### Data availability

Supplementary information, additional details for this study and R codes can be accessed at this link. Doi of the sequences submitted will be made available upon acceptance.

## Materials and methods

### Microbial origin

We isolated the microbes from wild *D. suzukii* flies that had infested fruit crops in the Netherlands (see table 1 for further details). Five females were crushed in 0.85% saline solution, diluted by the factor 10^-3^ to 10^-4^ and isolated on DYGS, R2A and YPDA medium by spread-plate method, followed by streak method. Isolated strains were grown overnight in liquid broth and stored in 20% sterilized glycerol at -80°C.

**Table 1.** List of microbial strains and combinations used for the assays.

| Strain identification | ID | Origin: flies infesting | Number of combinations as monocultures and cocultures |
| --- | --- | --- | --- |
| **Bacteria** |  |  | **3** |
| *Gluconobacter cerinus* PFA1 | GC1 | cherries in Randwijk, NL | Monocultures: GC1; GC2<br>Coculture: GC1+GC2 |
| *Gluconobacter cerinus* PJ7019 | GC2 | cherries in Randwijk, NL |  |
| **Yeast** |  |  | **7** |
| *Pichia kluyveri* PMM10-1742L | PK1 | cherries in Randwijk, NL | Monocultures: PK1; PK2; PT |
| *Pichia kluyveri*<br>SM12UFAM | PK2 | cherries in<br>Randwijk, NL | Cocultures:<br>PK1+PK2;PK2+PT;PT+PK1;PK1+<br>PK2+PT |
| *Pichia terricola* CBS 2617 | PT | strawberries in<br>Dirksland, NL |  |

### Microbial strain identification

To identify the isolated strains, we cloned the 16S rRNA gene for bacteria and ITS region for the yeasts into pGEM-T Easy Vector (USA Promega), which was inserted into JM109 competent cells according to the manufacturer’s protocol. We used the primers 5’ – AGAGTTTGTCMTGGCTCA-3’ and 5’-GGTTACCTTGTTACGACTT-3’ for 16S rRNA gene and 5’ – TCCGTA GGTGAACCTGCGG -3’ and 5’-TCCTCC GCTTATTGATATGC-3’ for ITS region. Sanger sequencing was performed at EUROFINS Biotech (Germany). Identification was performed using BioEdit (Hall, 2011) against the NCBI Blast (PCR conditions in the supplementary information). This study focused on two bacterial strains identified as *Gluconobacter cerinus*, named GC1 and GC2, and three yeast strains identified as belonging to the genus *Pichia*, PK1, PK2 and PT. We will subsequently use the abbreviations for these strains in the rest of the manuscript.

### Metabolic profiling

We assessed the carbon source utilization using Biolog plates – GenIII plates for bacteria and Yeast YT plates for yeasts – according to the manufacturer’s protocol (Biolog, CA, USA) and Puentez-Téllez and Salles (2018). Bacterial and yeast strains were grown overnight for ∼20 hours at 28°C and 180 rpm. Cells were then harvested and resuspended in saline solution and starved for 2.5-3 h. Next, bacterial and yeast cell suspensions were rinsed in sterile water and resuspended. For Biolog profiling, the final working absorbance was adjusted to OD590 0.03 for bacteria and OD590 0.9-1 for yeasts while adding them to the inoculation fluids. We added 100 µL of this solution to the wells and incubated them at 28°C for up to 70 h. Using a plate reader (Tecan Infinite 200, USA), we measured absorbance at 590 nm for bacteria and at 590 nm and 750 nm for yeasts (for additional details, see supplementary file, figure S1) at 13-time points from time point 0h until time point 70h according to the BIOLOG protocol. The background absorbance was removed by subtracting the blank OD values in the plates. We then used average well color development (AWCD) values for further characterization and analysis. We calculated AWCD values for all the wells at the final time point of 70 hours, using the formula: AWCD= ∑ Blank-corrected absorbance values of the wells/Total number of substrate sources

### Microbial strain preparation and addition to juvenile diet

To inoculate juvenile diets with microbial strains, we used overnight grown cultures and adjusted concentration to starting amounts of both bacterial and yeast cells of 10^5^ CFU/mL. A volume of 50 µL of these bacterial or yeast cell suspensions was added to fly food containing sucrose (40g/L), cornmeal (60g/L) and agar (7g/L). For monocultures, the total volume added per plate was 50 µL. Volume for cocultures of two strains was 25 µL/strain and for coculture of three strains was 16.66 µL/strain.

### Inoculation experiment

To assess the impact of microbial strains on juvenile development, we introduced eggs into plates with microbial mono or cocultures. First, we performed surface disinfection by submerging the eggs in 0.2% hypochlorite (ROTH, Germany) solution for 90 seconds. We then rinsed these eggs twice in sterile water and twice in saline and transferred the eggs into the food containing microbial treatments, with 13 eggs/plate. We used two control treatments: a negative control NYC (no yeast control) food, which had no yeast and where no to minimal sustenance of flies usually occurred, and a positive control HKY (heat-killed yeast) food which contained dead yeasts and would ensure high survival of the flies due to high nutritional conditions. The latter control treatment enabled us to assess the relative performance of the flies on the microbial strains compared to near-optimal conditions and to check the extent to which the bleach treatment affected the survival of the flies. Plates were then incubated at 20°C under a 16h/8h (light/dark) cycle, and larval development was monitored by daily observation. We used replicates of 36-41 plates for bacterial strains and 42-45 plates for yeast strains (Greiner dishes, size 35/10mm).

### Larval developmental assays

We calculated the <u>survival</u> by counting the number of pupae and the number of adults that emerged on the plates, including the live flies and dead adults that were partially stuck inside the food. We measured the <u>developmental time</u> of larvae, from larvae to pupae, and the developmental time from larvae to adult in the number of days. Furthermore, we collected the emerging flies and stored them in 70% ethanol for <u>morphometric measurements</u>. We measured the body size by taking the left hind tibia as the proxy, using an eye piece camera (Dino-Lite Digital Microscope, The Netherlands) attached to a microscope (Zeiss optical microscope) at a calibrated value of 0.5 mm scale. For bacterial treatment, we measured a subset of 29 flies. For yeast treatment, although we generated plots for all the strain combinations (with n=5 to 30), the statistical analysis was performed only for those with a minimal sample size of at least nine.

### Statistical analysis

All statistical analyses and figure generation was done in R version 4.0.3 (R core team, 2020), using the packages vegan (version 2.5-7; Oksanen et al., 2020), ggplot2 (version 3.3.3, Wickham, 2016), hlm2 (version 1.2.1, Marschner, 2011) and lme4 (version 1.1-26, Douglas et al., 2015). To assess the carbon use for metabolic characterization, we compared the functional groups in terms of average well color development (AWCD) using the Kruskal-Wallis test, followed by the posthoc Dunn test at a significance level of p-value of 0.05. We used binomial generalized linear models (GLMM) with binomial or poisson family to analyze the strains’ effect on fly survival and developmental time (as measured in days). For body measurement (tibia length), we used the mixed model ANOVA, incorporating both strain and sex as fixed effects and batch as a random effect. The significance level in all the analyses was at p-value of 0.05. For significant results, we further performed posthoc Tukey tests using the package emmeans (version 1.7.0, Searle et al., 1980) to identify the factors that contributed to the variation.

## Acknowledgement

This work was supported by the Adaptive Life Programme 2017 of the University of Groningen.

## Contribution

KG designed, performed experiments and statistical analyses, wrote the first draft, and edited subsequent drafts. JFS provided support with microbial culturing, metabolic profiling, and BW provided support with statistical analysis and data interpretation. JFS and BW provided intellectual input, commented on and edited subsequent drafts.

## Ethics approval

No approval was required for this study system.

## References

1. Anfora, G., Bouharoud, R., & Chebli, B. (2020). First report of Drosophila suzukii (Diptera: Drosophiladae) in North Africa. Moroccan Journal of Agricultural Sciences, 1(5).

2. Barber, A. E., Fleming, B. A., & Mulvey, M. A. (2016). Similarly, lethal strains of extraintestinal pathogenic Escherichia coli trigger markedly diverse host responses in a zebrafish model of sepsis. Msphere, 1(2), e00062–16.

3. Bing, X. L., Winkler, J., Gerlach, J., Loeb, G., & Buchon, N. (2021). Identification of natural pathogens from wild Drosophila suzukii. Pest management science, 77(4), 1594–1606.

4. Bing, X., Gerlach, J., Loeb, G., & Buchon, N. (2018). Nutrient-dependent impact of microbes on Drosophila suzukii development. MBio, 9(2), e02199–17.

5. Braga, R. M., Dourado, M. N., & Araújo, W. L. (2016). Microbial interactions: ecology in a molecular perspective. Brazilian Journal of Microbiology, 47, 86–98.

6. Chown, S. L., & Terblanche, J. S. (2006). Physiological diversity in insects: ecological and evolutionary contexts. Advances in insect physiology, 33, 50–152.

7. Crotti, E., Balloi, A., Hamdi, C., Sansonno, L., Marzorati, M., Gonella, E., Favia, G., Cherif, A., Bandi, C., Alma, A. and Daffonchio, D. (2012), Microbial symbionts: a resource for the management of insect-related problems. Microbial Biotechnology, 5: 307–317.

8. Crotti, E., Rizzi, A., Chouaia, B., Ricci, I., Favia, G., Alma, A., … & Daffonchio, D. (2010). Acetic acid bacteria, newly emerging symbionts of insects. Applied and Environmental Microbiology, 76(21), 6963–6970.

9. Datasheet. (2021) Drosophila suzukii report

10. De Muynck, C., Pereira, C. S., Naessens, M., Parmentier, S., Soetaert, W., & Vandamme, E. J. (2007). The genus Gluconobacter oxydans: comprehensive overview of biochemistry and biotechnological applications. Critical reviews in biotechnology, 27(3), 147–171.

11. Deppenmeier, U., & Ehrenreich, A. (2009). Physiology of acetic acid bacteria in light of the genome sequence of Gluconobacter oxydans. Journal of molecular microbiology and biotechnology, 16(1-2), 69–80.

12. Douglas Bates, Martin Maechler, Ben Bolker, Steve Walker (2015). Fitting Linear Mixed-Effects Models Using lme4. Journal of Statistical Software, 67(1), 1–48.

13. Douglas, A. E. (2018). Contradictory results in microbiome science exemplified by recent Drosophila research. MBio, 9(5), e01758–18.

14. Dwivedi, M. (2020). Gluconobacter. In Beneficial Microbes in Agro-Ecology (pp. 521–544). Academic Press.

15. Early, A. M., Shanmugarajah, N., Buchon, N., & Clark, A. G. (2017). Drosophila genotype influences commensal bacterial levels. PloS one, 12(1), e0170332.

16. Fountain, M.T., Bennett, J., Cobo-Medina, M., Conde Ruiz, R., Deakin, G., Delgado, A., Harrison, R. and Harrison, N. (2018). Alimentary microbes of winter-form Drosophila suzukii. Insect molecular biology, 27(3), 383–392.

17. Gasmi, L., Baek, S., Kim, J.C., Kim, S., Lee, M.R., Park, S.E., Shin, T.Y., Lee, S.J., Parker, B.L. and Kim, J.S. (2021). Gene diversity explains variation in biological features of insect killing fungus, Beauveria bassiana. Scientific reports, 11(1), 1–12.

18. Gould, A. L., Zhang, V., Lamberti, L., Jones, E. W., Obadia, B., Korasidis, N., … & Ludington, W. B. (2018). Microbiome interactions shape host fitness. Proceedings of the National Academy of Sciences, 115(51), E11951–E11960.

19. H. Wickham. ggplot2: Elegant Graphics for Data Analysis. Springer-Verlag, New York, 2016.

20. Hall, T., Biosciences, I., & Carlsbad, C. (2011). BioEdit: an important software for molecular biology. GERF Bull Biosci, 2(1), 60–61.

21. Hamby, K. A., Hernández, A., Boundy-Mills, K., & Zalom, F. G. (2012). Associations of yeasts with spotted-wing Drosophila (Drosophila suzukii; Diptera: Drosophilidae) in cherries and raspberries. Applied and environmental microbiology, 78(14), 4869–4873.

22. Hassani, M., Durán, P., & Hacquard, S. (2018). Microbial interactions within the plant holobiont. Microbiome, 6(1), 1–17.

23. Hauser, M. (2011). A historic account of the invasion of Drosophila suzukii (Matsumura)(Diptera: Drosophilidae) in the continental United States, with remarks on their identification. Pest management science, 67(11), 1352–1357.

24. He, M., Jiang, J., & Cheng, D. (2017). The plant pathogen Gluconobacter cerinus strain CDF1 is beneficial to the fruit fly Bactrocera dorsalis. Amb Express, 7(1), 1–10.

25. Hu, C., Rio, R.V., Medlock, J., Haines, L.R., Nayduch, D., Savage, A.F., Guz, N., Attardo, G.M., Pearson, T.W., Galvani, A.P. and Aksoy, S. (2008). Infections with immunogenic trypanosomes reduce tsetse reproductive fitness: potential impact of different parasite strains on vector population structure. PLoS neglected tropical diseases, 2(3), e192.

26. Itoh, H., Jang, S., Takeshita, K., Ohbayashi, T., Ohnishi, N., Meng, X.Y., Mitani, Y. and Kikuchi, Y. (2019). Host–symbiont specificity determined by microbe–microbe competition in an insect gut. Proceedings of the National Academy of Sciences, 116(45), 22673–22682.

27. Jakob, F., Quintero, Y., Musacchio, A., Estrada-de los Santos, P., Hernández, L., & Vogel, R. F. (2019). Acetic acid bacteria encode two levansucrase types of different ecological relationship. Environmental microbiology, 21(11), 4151–4165.

28. Jari Oksanen, F. Guillaume Blanchet, Michael Friendly, Roeland Kindt, Pierre Legendre, Dan McGlinn, Peter R. Minchin, R. B. O’Hara, Gavin L. Simpson, Peter Solymos, M. Henry H. Stevens, Eduard Szoecs and Helene Wagner (2020). vegan: Community Ecology Package. R package version 2.5–7. https://CRAN.R-project.org/package=vegan

29. Joop, G., & Vilcinskas, A. (2016). Coevolution of parasitic fungi and insect hosts. Zoology, 119(4), 350–358.

30. Katzen, F., Ferreiro, D. U., Oddo, C. G., Ielmini, M. V., Becker, A., Pühler, A., & Ielpi, L. (1998). Xanthomonas campestris pv. campestris gum mutants: effects on xanthan biosynthesis and plant virulence. Journal of bacteriology, 180(7), 1607–1617.

31. Kenis, M., Tonina, L., Eschen, R., van der Sluis, B., Sancassani, M., Mori, N., … & Helsen, H. (2016). Non-crop plants used as hosts by Drosophila suzukii in Europe. Journal of Pest Science, 89(3), 735–748.

32. Kinnula, H., Mappes, J., & Sundberg, L. R. (2017). Co-infection outcome in an opportunistic pathogen depends on the inter-strain interactions. BMC evolutionary biology, 17(1), 1–10.

33. Kroll, S., Agler, M. T., & Kemen, E. (2017). Genomic dissection of host–microbe and microbe–microbe interactions for advanced plant breeding. Current Opinion in Plant Biology, 36, 71–78.

34. Kwadha, C. A., Okwaro, L. A., Kleman, I., Rehermann, G., Revadi, S., Ndlela, S., … & Becher, P. G. (2021). Detection of the spotted wing drosophila, Drosophila suzukii, in continental sub-Saharan Africa. Journal of Pest Science, 94(2), 251–259.

35. Lasa, R., Navarro-de-la-Fuente, L., Gschaedler-Mathis, A. C., Kirchmayr, M. R., & Williams, T. (2019). Yeast species, strains, and growth media mediate attraction of Drosophila suzukii (Diptera: Drosophilidae). Insects, 10(8), 228.

36. Lee, H. Y., Lee, S. H., Lee, J. H., Lee, W. J., & Min, K. J. (2019). The role of commensal microbes in the lifespan of Drosophila melanogaster. Aging (Albany NY), 11(13), 4611.

37. Lewis, M. T., & Hamby, K. A. (2019). Differential impacts of yeasts on feeding behavior and development in larval Drosophila suzukii (Diptera: Drosophilidae). Scientific reports, 9(1), 1–12.

38. Liu, R., Zhang, Q., Chen, F., & Zhang, X. (2015). Analysis of culturable yeast diversity in spontaneously fermented orange wine, orange peel and orangery soil of a Ponkan plantation in China. Annals of microbiology, 65(4), 2387–2391.

39. Lynch, K. M., Zannini, E., Wilkinson, S., Daenen, L., & Arendt, E. K. (2019). Physiology of acetic acid bacteria and their role in vinegar and fermented beverages. Comprehensive Reviews in Food Science and Food Safety, 18(3), 587–625.

40. Marschner, I. C. (2011). glm2: Fitting generalized linear models with convergence problems. The R Journal 3(2): 12–15

41. Matsushita, K., Fujii, Y., Ano, Y., Toyama, H., Shinjoh, M., Tomiyama, N., Miyazaki, T., Sugisawa, T., Hoshino, T. and Adachi, O., (2003). 5-Keto-d-gluconate production is catalyzed by a quinoprotein glycerol dehydrogenase, major polyol dehydrogenase, in Gluconobacter species. Applied and Environmental Microbiology, 69(4), 1959–1966.

42. Mazzetto, F., Gonella, E., Crotti, E., Vacchini, V., Syrpas, M., Pontini, M., Mangelinckx, S., Daffonchio, D. and Alma, A. (2016). Olfactory attraction of Drosophila suzukii by symbiotic acetic acid bacteria. Journal of Pest Science, 89(3), 783–792.

43. McLean, A. H. C., Hrček, J., Parker, B. J., Mathé-Hubert, H., Kaech, H., Paine, C., & Godfray, H. C. J. (2020). Multiple phenotypes conferred by a single insect symbiont are independent. Proceedings of the Royal Society B, 287(1929), 20200562.

44. McLean, A. H., Parker, B. J., Hrček, J., Kavanagh, J. C., Wellham, P. A., & Godfray, H. C. J. (2018). Consequences of symbiont co-infections for insect host phenotypes. Journal of Animal Ecology, 87(2), 478–488.

45. Miraldo, A., & Duplouy, A. (2019). High Wolbachia strain diversity in a clade of dung beetles endemic to Madagascar. Frontiers in ecology and evolution, 7, 157.

46. Narita, S., Nomura, M., & Kageyama, D. (2007). Naturally occurring single and double infection with Wolbachia strains in the butterfly Eurema hecabe: transmission efficiencies and population density dynamics of each Wolbachia strain. FEMS microbiology ecology, 61(2), 235–245.

47. Niu, B., Wang, W., Yuan, Z., Sederoff, R. R., Sederoff, H., Chiang, V. L., & Borriss, R. (2020). Microbial interactions within multiple-strain biological control agents impact soil-borne plant disease. Frontiers in Microbiology, 2452.

48. Pang, R., Chen, M., Yue, L., Xing, K., Li, T., Kang, K., Liang, Z., Yuan, L. and Zhang, W. (2018). A distinct strain of Arsenophonus symbiont decreases insecticide resistance in its insect host. PLoS genetics, 14(10), e1007725.

49. Puentes-Téllez, P. E., & Falcao Salles, J. (2018). Construction of effective minimal active microbial consortia for lignocellulose degradation. Microbial ecology, 76(2), 419–429.

50. R Core Team (2020). R: A language and environment for statistical computing. R Foundation for Statistical Computing, Vienna, Austria. URL: https://www.R-project.org/.

51. Raspor, P., & Goranovič, D. (2008). Biotechnological applications of acetic acid bacteria. Critical reviews in biotechnology, 28(2), 101–124.

52. Russel, J., Røder, H. L., Madsen, J. S., Burmølle, M., & Sørensen, S. J. (2017). Antagonism correlates with metabolic similarity in diverse bacteria. Proceedings of the National Academy of Sciences, 114(40), 10684–10688.

53. Sainz, F., Mas, A., & Torija, M. J. (2016). Draft genome sequences of Gluconobacter cerinus CECT 9110 and Gluconobacter japonicus CECT 8443, acetic acid bacteria Isolated from grape must. Genome announcements, 4(3), e00621–16.

54. Searle, S. R., Speed, F. M., & Milliken, G. A. (1980). Population marginal means in the linear model: an alternative to least squares means. The American Statistician, 34(4), 216–221.

55. Seth, E. C., & Taga, M. E. (2014). Nutrient cross-feeding in the microbial world. Frontiers in microbiology, 5, 350.

56. Shelly, T. E. (2018). Larval host plant influences male body size and mating success in a tephritid fruit fly. Entomologia Experimentalis et Applicata, 166(1), 41–52.

57. Smee, M. R., Baltrus, D. A., & Hendry, T. A. (2017). Entomopathogenicity to two hemipteran insects is common but variable across epiphytic Pseudomonas syringae strains. Frontiers in Plant Science, 8, 2149.

58. Sofo, A., & Ricciuti, P. (2019). A standardized method for estimating the functional diversity of soil bacterial community by biolog® EcoPlatesTM assay—The case study of a sustainable olive orchard. Applied Sciences, 9(19), 4035.

59. Solomon, G. M., Dodangoda, H., McCarthy-Walker, T., Ntim-Gyakari, R., & Newell, P. D. (2019). The microbiota of Drosophila suzukii influences the larval development of Drosophila melanogaster. PeerJ, 7, e8097.

60. Sombolestani, A. S., Cleenwerck, I., Cnockaert, M., Borremans, W., Wieme, A. D., De Vuyst, L., & Vandamme, P. (2021). Characterization of novel Gluconobacter species from fruits and fermented food products: Gluconobacter cadivus sp. nov., Gluconobacter vitians sp. nov. and Gluconobacter potus sp. nov. International Journal of Systematic and Evolutionary Microbiology, 71(3), 004751.

61. Spitaler, U., Bianchi, F., Eisenstecken, D., Castellan, I., Angeli, S., Dordevic, N., … & Schmidt, S. (2020). Yeast species affects feeding and fitness of Drosophila suzukii adults. Journal of Pest Science, 93(4), 1295–1309.

62. Stefanini, I. (2018). Yeast-insect associations: It takes guts. Yeast, 35(4), 315–330.

63. Sugio, A., Dubreuil, G., Giron, D., & Simon, J. C. (2015). Plant–insect interactions under bacterial influence: ecological implications and underlying mechanisms. Journal of experimental botany, 66(2), 467–478.

64. Tahara, Y., Yamada, Y., & Kondo, K. (1976). Phospholipid composition of Gluconobacter cerinus. Agricultural and Biological Chemistry, 40(12), 2355–2360.

65. Tran, J., Aksenov, V., & Rollo, C. D. (2018). A multi-ingredient athletic supplement disproportionately enhances hind leg musculature, jumping performance, and spontaneous locomotion in crickets (A cheta domesticus). Entomologia Experimentalis et Applicata, 166(1), 63–73.

66. Vacchini, V., Gonella, E., Crotti, E., Prosdocimi, E. M., Mazzetto, F., Chouaia, B., … & Daffonchio, D. (2017). Bacterial diversity shift determined by different diets in the gut of the spotted wing fly Drosophila suzukii is primarily reflected on acetic acid bacteria. Environmental microbiology reports, 9(2), 91–103.

67. Van Keer, C., Abeele, P. V., Swings, J., Gossele, F., & De Ley, J. (1981). Acetic acid bacteria as causal agents of browning and rot of apples and pears. Zentralblatt für Bakteriologie Mikrobiologie und Hygiene: I. Abt. Originale C: Allgemeine, angewandte und ökologische Mikrobiologie, 2(2), 197–204.

68. Van Rossum, T., Ferretti, P., Maistrenko, O.M. et al. Diversity within species: interpreting strains in microbiomes. Nat Rev Microbiol 18, 491–506 (2020).

69. Venturelli, O.S., Carr, A.V., Fisher, G., Hsu, R.H., Lau, R., Bowen, B.P., Hromada, S., Northen, T. and Arkin, A.P. (2018). Deciphering microbial interactions in synthetic human gut microbiome communities. Molecular systems biology, 14(6), e8157.

70. Vesga, P., Augustiny, E., Keel, C., Maurhofer, M., & Vacheron, J. (2021). Phylogenetically closely related pseudomonads isolated from arthropods exhibit differential insect-killing abilities and genetic variations in insecticidal factors. Environmental Microbiology.

71. Vicente, J., Calderón, F., Santos, A., Marquina, D., & Benito, S. (2021). High potential of Pichia kluyveri and other Pichia species in wine technology. International Journal of Molecular Sciences, 22(3), 1196.

72. Vicente, J., Calderón, F., Santos, A., Marquina, D., & Benito, S. (2021). High potential of Pichia kluyveri and other Pichia species in wine technology. International Journal of Molecular Sciences, 22(3), 1196.

73. Walsh, D. B., Bolda, M. P., Goodhue, R. E., Dreves, A. J., Lee, J., Bruck, D. J., … & Zalom, F. G. (2011). Drosophila suzukii (Diptera: Drosophilidae): invasive pest of ripening soft fruit expanding its geographic range and damage potential. Journal of Integrated Pest Management, 2(1), G1–G7.

74. Wieme, A. D., Spitaels, F., Aerts, M., De Bruyne, K., Van Landschoot, A., & Vandamme, P. (2014). Identification of beer-spoilage bacteria using matrix-assisted laser desorption/ionization time-of-flight mass spectrometry. International journal of food microbiology, 185, 41–50.

75. Wilcoxon, F. (1945). Individual comparisons by ranking methods. Biom. Bull., 1, 80– 83.

76. Xie, S., Lan, Y., Sun, C., & Shao, Y. (2019). Insect microbial symbionts as a novel source for biotechnology. World Journal of Microbiology and Biotechnology, 35(2), 25.

77. Yan, Y., Nguyen, L. H., Franzosa, E. A., & Huttenhower, C. (2020). Strain-level epidemiology of microbial communities and the human microbiome. Genome medicine, 12(1), 1–16.

