## Supplementary file for "Insights into the microbial strain mediated impact on pest insect development"

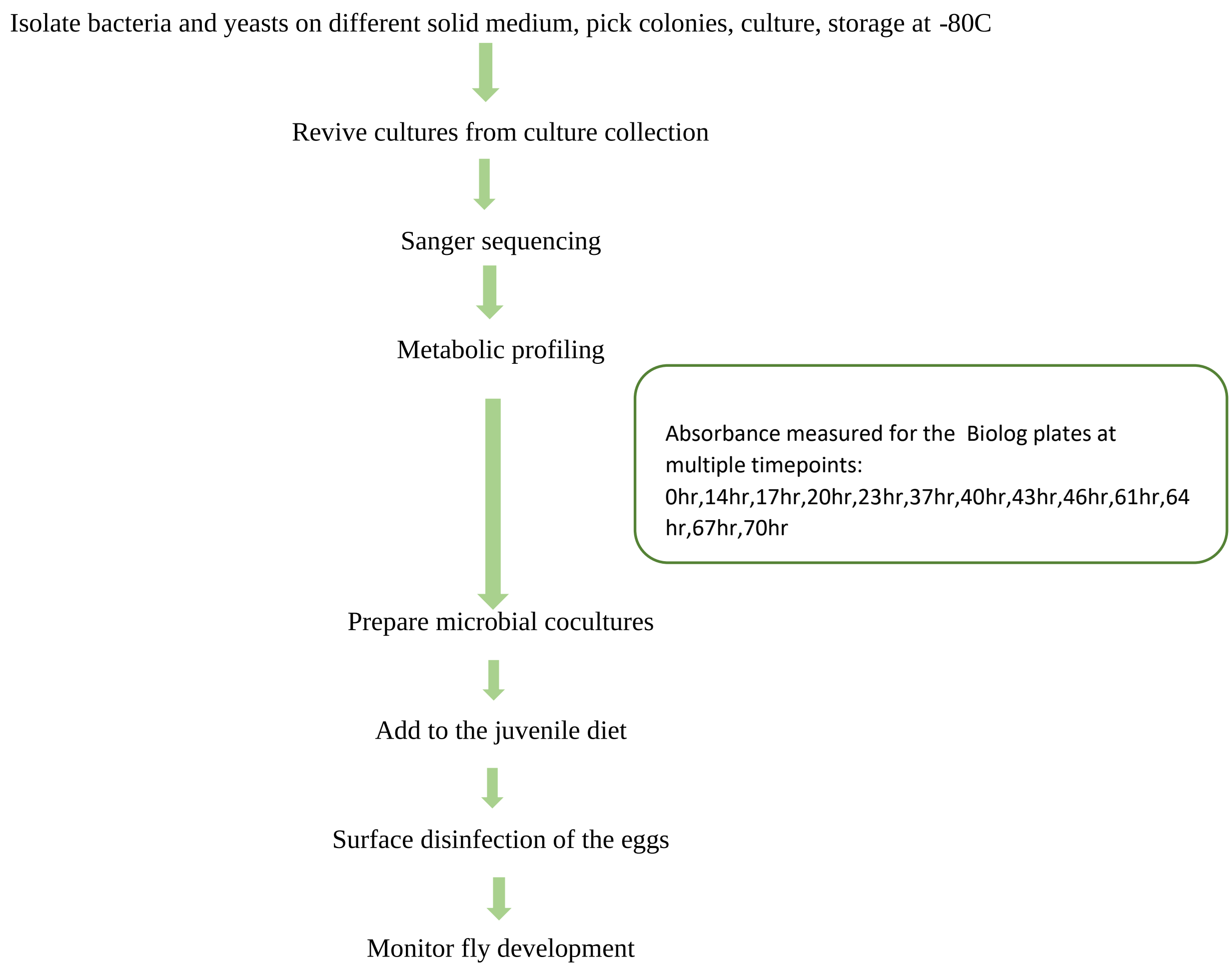


[note: liquid cultures when used in this study were incubated at 28 degrees Celsius and 180 rpm in shaker incubator]

**Figure S1: Outline of the complete experimental set-up used in this study.**

**Culture medium used for bacteria and yeasts isolation and revival**

**DYGS medium**

d$H_2O$ – 1L

glucose - 2 g

yeast extract - 2 g

peptone - 1.5 g

glutamic acid - 1.3 g

Dipotassium phosphate - 500 mg

Epsom salt - 500 mg

pH - 6.0

Agar 12g

**Bacteria: Mannitol broth**

Peptic digest – 2.8g/L

Yeast extract-5g/L

Mannitol-25g/L

**Yeasts: Potato dextrose broth (premade; Oxoid, Thermo Fisher Scientific)**

**Primers used in Sanger sequencing**

Bacteria:16S rRNA gene

Forward: AGAGTTTGTCMTGGCTCAG

Reverse: GGTTACCTTGTTACGACTT

Yeast: ITS region

Forward: TCCGTA GGTGAACCTGCGG

Reverse: TCCTCC GCTTATTGATATGC

**PCR conditions for amplifying the target region**

94C – 3min

94C – 1min

53C – 1min **35 cycles**

72C – 1min

72C – 4min

| Colony id | Abbreviation used in this study | % query cover | Closest match in NCBI | Accession | % identity |
|---|---|---|---|---|---|
| D5047 | GC1 | 99 | *Gluconobacter cerinus* PFA1 | KP234004.1 | 99.46 |
| R504 | GC2 | 99 | *Gluconobacter cerinus* PJ7019 | MG266178.1 | 98.70 |
| Y502 | PK1 | 86 | *Pichia kluyveri* PMM10-1742L | MN268784.1 | 93.82 |
| Y5042 | PK2 | 85 | *Pichia kluyveri* SM12UFAM | KP132503.1 | 99.57 |
| YS3 | PT | 84 | *Pichia terricola* CBS 2617 | NR153294.1 | 99.77 |

**Supplementary table: Details of the closest hits identified using NCBI Blast tool for the bacterial and yeast strains**
